# A Systematic Benchmark of High-Accuracy PacBio Long-Read RNA Sequencing for Transcript-Level Quantification

**DOI:** 10.1101/2025.05.30.656561

**Authors:** David Wissel, Madison M. Mehlferber, Khue M. Nguyen, Vasilii Pavelko, Elizabeth Tseng, Mark D. Robinson, Gloria M. Sheynkman

## Abstract

PacBio long-read RNA sequencing resolves transcripts with greater clarity than short-read technologies, yet its quantitative performance remains under-evaluated at scale. Here, we benchmark the high-throughput PacBio Kinnex platform against Illumina short-read RNA-seq using matched, deeply sequenced datasets across a time course of endothelial cell differentiation. Compared to Illumina, Kinnex achieved comparable gene-level quantification and more accurate transcript discovery and transcript quantification. While Illumina detected more transcripts overall, many reflected potentially unstable or ambiguous estimates in complex genes. Kinnex largely avoids these issues, producing more reliable differential transcript expression (DTE) calls, despite a mild bias against short transcripts (shorter than 1.25 kb). When correcting Illumina for inferential variability, Kinnex and Illumina quantifications were highly concordant, demonstrating equivalent performance. We also benchmarked long-read tools, nominating Oarfish as the most efficient for our Kinnex data. Together, our results establish Kinnex as a reliable platform for full-length transcript quantification.

## Introduction

RNA-sequencing (RNA-seq) has enabled comprehensive measurements of the transcriptome, but standard short-read approaches face inherent limitations in resolving full-length transcripts. The assembly of fragmented reads into complete transcripts is error-prone, particularly for genes with complex splicing, resulting in ambiguity in transcript discovery and quantification [1]. PacBio’s long-read RNA-seq (lrRNA-seq) platform addresses this limitation by sequencing entire RNA molecules. However, until recently, the low throughput (less than 10 M reads per sample) limited its utility for quantitative analyses, a finding confirmed by bench-marking efforts such as the Long-Read RNA-seq Genome annotation Assessment Consortium (LRGASP) [2].

The MAS-Iso-seq method, commercialized as PacBio Kinnex, employs a cDNA concatenation approach that increases read yield on average by 8-fold relative to previous protocols [3]. This throughput improvement could enable new experimental designs that combine the transcript resolution of long reads with the quantitative depth required for transcriptome profiling. Nonetheless, a rigorous evaluation of Kinnex for transcript quantification has not yet been performed. Previous studies have focused primarily on earlier PacBio chemistries [4, 5] or have emphasized the potential for transcript discovery rather than quantification [6, 7]. Most exisiting benchmarking studies for long-read quantification more broadly have focused on Oxford Nanopore (ONT) data, which remains more error-prone than PacBio [5, 6]. As a result, it remains unclear how Kinnex compares to Illumina across complex biological samples, which tools best quantify its transcript abundances, and how it performs in differential expression analyses.

In this study, we evaluate the performance of PacBio Kinnex long-read RNA-seq for transcript-level quantification, benchmarked against standard Illumina short-read RNA-seq. Using matched, deeply sequenced datasets collected across a five-day time course of human iPSC differentiation to endothelial cells, we systematically compare transcript discovery, abundance quantification, and downstream tasks such as differential gene and transcript expression. We also assess the performance of multiple long-read quantification tools and analyze sources of quantification discordance between platforms. Together, our results help to define the capabilities and limitations of Kinnex for transcript-resolved RNA-seq at scale.

## Results

### Kinnex lrRNA-seq Is Technically Robust and Enables Effective Biological Downstream Analysis

We collected RNA from human induced pluripotent stem cells (iPSCs) (WTC11) cells undergoing a five-day endothelial differentiation time course [8], and performed matched sequencing using both Kinnex (PacBio Revio) and Illumina (NovaSeq) platforms (see **Methods**). To assess technical performance, we spiked in Spike-In RNA variants (SIRV) controls (Figure 1A). For each day of differentiation, we isolated RNA with three replicates for day zero and five, two replicates for day three, and one replicate for the remaining days (one, two, and four) (Figure 1A). We excluded the single day-two replicate from further analysis due to quality issues (Figure S1A-J). Illumina libraries yielded approximately 67 million short reads per sample on average (Figure 1B). In parallel, Kinnex produced approximately 54 million full-length reads per sample, on average, with a median length of ~1.55-kb—roughly 5.2 times longer than the 299-bp paired-end Illumina reads, and thus a higher total number of bases sequenced per sample (Figure 1C-D). Both technologies achieved high alignment rates to the human genome (Illumina: ~95%, Kinnex: ~97%; Figure 1B), along with high average base quality scores (Illumina: Phred~36, Kinnex: Phred~38; Figure 1F). In terms of empirical error profiles, Kinnex showed more indel-related errors compared to Illumina (Kinnex average edit distance (indels): ~0.00205, Illumina average edit distance (indels): ~0.0000004), with SNVs showing the reverse behavior (Kinnex average edit distance (SNVs): ~0.00087, Illumina average edit distance (SNVs): ~0.00301) (Figure 1G). Both technologies showed a slight 3*^′^* bias for shorter transcripts (shorter than one kb), with this bias improving for transcripts of medium length with increasing length (between one and two kb, two and three kb, and three and six kb) (Figure 1H). For long transcripts (longer than six kb), Illumina exhibited a moderate 5*^′^* bias, whereas Kinnex showed a U-shaped coverage, with distinctly lower coverage around the middle of the gene body. We note that the behavior of long-reads not fully covering very long transcripts is consistent with previous works and may be attributable to a combination of internal priming and the difficulty of RT for long transcripts [9, 10].

**Figure 1:**
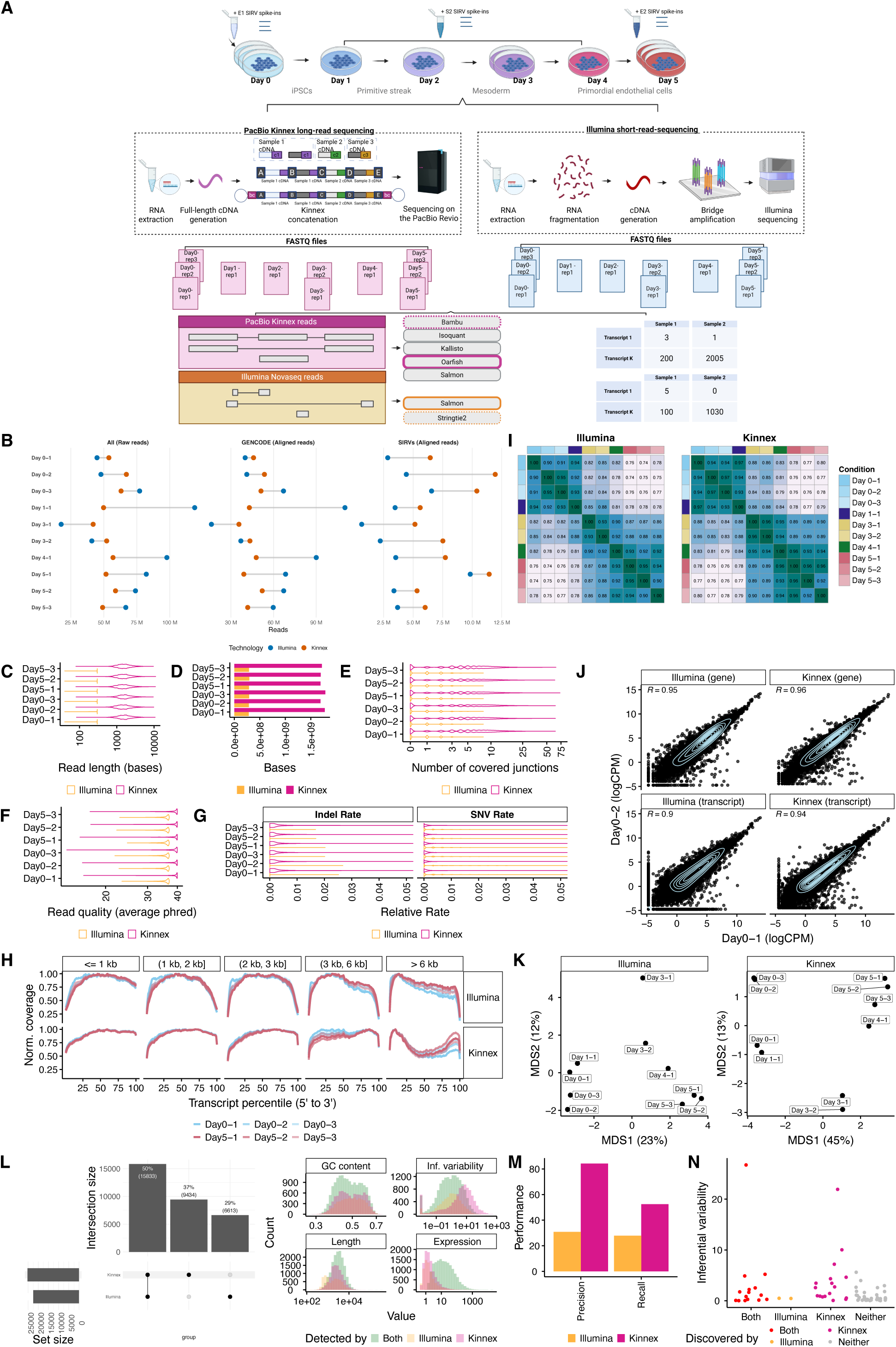
Dataset quality control, summary statistics and characterization. **A.** Experimental design to collect a matched RNA-sequencing dataset of differentiated primordial endothelial cells to track transcript dynamics over time. The Kinnex protocol emphasizes the use of full-length cDNA, distinguishing it from short-read RNA sequencing sample preparation workflows. Kinnex preparation abbreviations (bc1 = barcode 1, bc2 = barcode 2, A, B, C and D denote Kinnex-specific primers, c1, c2 and c3 represent individual cDNA molecules. Respective transcript quantification and discovery methods employed for each technology are denoted. **B.** Number of raw and aligned reads per technology, stratified by their source across all samples. **C.** Read lengths of 1M randomly sampled reads aligning to the human genome, stratified by technology for Days 0 and 5. We highlight Days 0 and 5 due to their replication. For Illumina, the length was calculated across both ends. **D.** Number of base pairs sequenced from 1M randomly sampled reads collected for each technology (same reads as **C**). **E.** Number of covered junctions per read from 1M randomly sampled reads collected for each technology (same reads as **C**). **F.** Average base quality of 1M randomly sampled reads aligning to the human genome, stratified by technology for Days 0 and 5 (same reads as **C**). The average base quality for Illumina was determined from both ends. **G.** Average edit distance per read, stratified by indel vs SNV errors, of 1M randomly sampled reads collected for each technology (same reads as **C**).**H.** Normalized coverage of reads aligning to 2,500 randomly sampled GENCODE transcripts each across gene body percentiles, stratified by technology and transcript length, across all samples. **I.** Spearman correlation heatmap of transcript-level quantifications highlighting sample similarities across the full differentiation for each technology. **J.** Scatter plot for Day 0-1 and Day 0-2 replicates at the gene and transcript levels following quantification, stratified by technology. **K.** Transcript-level quantification-based MDS plots highlighting sample differences across the full differentiation, stratified by technology. **L.** Left: Upset plot of the transcripts detected on Day 0 (>1 CPM in all three replicates) for both technologies. Right: Attributes of transcripts detected by both or only one of the technologies. GC content, Inferential variability as measured by the inferential relative variance on Day 0 Illumina samples, transcript length, and median expression in both of the technologies. **M.** Precision and recall of de-novo transcript discovery on three Day 0 (E1 mix) SIRV replicates by technology. Both platforms were ran on downsampled read sets consisting of 2.5 M reads. **N.** Mean inferential variability of SIRV transcripts on three Day 0 replicates by whether they were discovered by both platforms, only Illumina, only Kinnex, or neither platform. Figure 1A was created in Biorender, Mehlferber, M. (2025) Biorender.

Next, we leveraged inferential variability, a measure of the degree of uncertainty when estimating the abundance of a specific transcript. This uncertainty may become a major issue when short reads map ambiguously to multiple transcripts (see **Methods**). Using Salmon for the quantification of Illumina samples and Oarfish for the quantification of Kinnex samples, and adjusting for the inferential variability in Illumina samples using existing approaches [11], we observed decreasing correlations between transcript-level sample expression over the course of the differentiation time-course as well as high reproducibility of gene and transcript quantification for replicates of the same day (Spearman > 0.9) (Figure 1I-J). Multidimensional scaling (MDS) and Spearman correlation analyses showed tight clustering among same-day replicates and a clear separation along the differentiation axis for both platforms (Figure 1J-K). At full depth, on Day 0, Kinnex detected a total of 25,267 transcripts (where detection was defined by an expression of 1 CPM or greater in all three replicates), 2,821 more compared to Illumina (Figure 1L, left). The pattern of Kinnex detecting more transcripts than Illumina also held when defining detection using a simple read count threshold at lower, downsampled depths (Supplementary Table S1). The two main patterns in terms of which technology tended to detect which transcripts were the following: First, Illumina detected more transcripts that were relatively short (1 kb - 1.25 kb) compared to Kinnex (Figure 1, right). However, Illumina struggled with the detection of transcripts that exhibited high inferential variability (Figure 1, right). In terms of expression and GC content, there were no clear patterns as far as differences in detection between the technologies went.

Differential Transcript Usage (DTU) analysis revealed prominent transcript switches across the differentiation, including within transcripts of the *IPO11* gene (Figure S2A, Supplementary Table S2). Marker gene expression matched biological expectations associated with the differentiation, including *NANOG* downregulation and endothelial cell (EC) marker induction (*KDR*, *ETV2*, *CD34*, *SOX7*, and *SOX17*) (Figure S2B and Supplementary Table S3) [12–16].

Collectively, these results demonstrate that Kinnex reads are comparable in quality to those from Illumina, are able to capture expected biological patterns, and are suitable for a technical comparison.

### Kinnex Enables the Discovery of Novel Transcripts

We next assessed the potential for Kinnex and Illumina to discover transcripts (also known as transcript identification). We used the SIRV spike-ins, which provide a known set of sequences introduced during library preparation.

Using Bambu for the discovery of novel transcripts in Kinnex samples and StringTie2 for Illumina samples, both with default parameters, we found that Kinnex outperformed Illumina in both precision and recall, indicating more complete and accurate transcript discovery when using long-read data (Figure 1M). To understand why Illumina-based transcript discovery was incomplete, we examined transcripts that were identified by Kinnex but missed by Illumina. We found that such transcripts tended to have more complex splicing patterns, as evidenced by higher inferential variability—such as a large number of alternative exons—which are challenging to resolve with short reads, although the pattern was not fully clear (Figure 1N).

Given Kinnex’s greater reliability for transcript discovery, we used it as the basis for identifying novel transcripts in the differentiation samples. We ran Bambu on the Kinnex-aligned reads and classified the transcripts using SQANTI3, leading to the identification of 12,341 novel transcripts and 515 novel genes (Figure S2C; Supplementary Table S4) [17]. Of the 12,341 novel transcripts identified using Bambu and classified with SQANTI3, the majority (~89%) were lowly expressed (less than 10 CPM). However, 1,337 transcripts (10.8%) were at least moderately expressed (10 CPM or greater) and multi-exonic (SQANTI3 categories: ISM, NIC, or NNC; Figure S2C) [2, 18]. Among these, 1,097 (8.9% of all novel transcripts) were predicted to be protein-coding based on ORFanage. Overall, 77.5% (9,565/12,341) of all novel transcripts were predicted to be protein-coding. As an illustrative example, BambuTx5158 is a novel transcript with moderate expression (34 CPM), overlapping the *RP11* gene locus (ENSG00000254202), that has strong read support from both Illumina and Kinnex (Figure S2D), but is predicted to be non-coding.

### Benchmarking of Kinnex Quantification Tools Shows Strong Performance of Oarfish

Although Kinnex long-read sequencing excelled at discovering novel transcripts, its ability to measure the abundances of transcripts remained uncertain. Therefore, we next benchmarked the performance of Kinnex quantification tasks across several facets of transcriptome quantification, in comparison to Illumina.

To perform an in-depth characterization of transcript quantification, we first attempted to determine suitable quantification tools for Kinnex data, as long-read analysis methods are still in the process of refinement, while methods for the quantification of short-read RNA-seq (srRNA-seq) are comparatively well established [19]. We conducted a systematic comparison of five leading tools—Oarfish, lr-kallisto, Isoquant, Bambu, and Salmon—using read-depth-matched Kinnex datasets and SIRV spike-ins as ground truth [17, 20–23].

First, across the five tools, we compared the computational resource usage requirements needed by running different numbers of subsampled reads on the same hardware. In these efficiency evaluations, Oarfish and lr-kallisto stood out for their fast runtime and low memory usage across read depths (Figure 2A).

**Figure 2:**
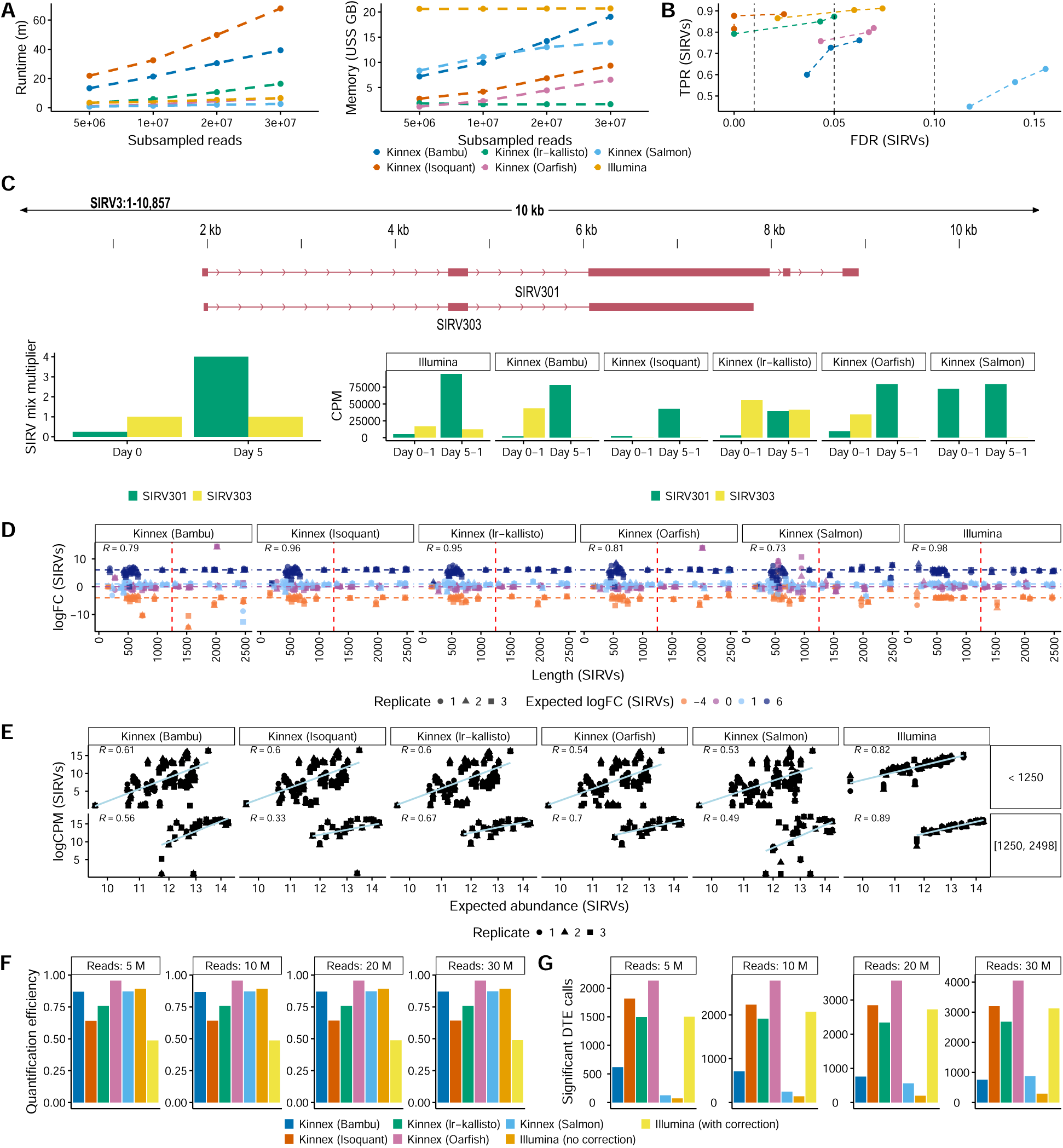
Comparison between different Kinnex quantification methods. **A.** Computational requirements of quantification methods for Kinnex and Illumina in terms of runtime (left) and memory (right) across all days and replicates of differentiation after downsampling the GENCODE-aligned data of each day to a fixed read depth (see **Methods**). Alignment and index creation were not included within computational requirements. **B.** True positive rates (TPRs) and false discovery rates (FDRs) for Kinnex and Illumina quantification methods on Differential Transcript Expression (DTE) between Day 0 (E1 mix) and Day 5 (E2 mix) on the SIRV spike-ins when downsampling to a fixed depth of 1M reads for all samples (see **Methods**). **C.** Top: Browser track highlighting an example of a pair of problematic SIRV transcripts that may cause outliers for some Kinnex lrRNA-seq quantification methods. Bottom: Highlight of how the SIRV301 and SIRV303 SIRV transcript changes between the mixes utilized for Day 0 and Day 5 samples. While the SIRV303 transcript stays constant in expression between mixes, the SIRV301 transcript increases its expression drastically (16-fold), leading to most of the Kinnex methods attributing all expression to the SIRV301 sample in the Day 5 mix, since the SIRV303 transcript is fully contained within the SIRV301 transcript. **D.** Relative quantification results between the same replicate on Day 0 (E1 mix) versus Day 5 (E2 mix) of four quantification methods for Kinnex lrRNA-seq and Illumina as measured by similarity of observed and expected log-fold changes between SIRV spike-in mixes across SIRV transcript length when downsampling to a fixed depth of 1M reads for all samples (see **Methods**). **E.** Absolute quantification of four quantification methods for Kinnex lrRNA-seq and Illumina as measured by concordance of observed and expected absolute abundances for each replicate on Day 0 (E1 mix) when downsampling to a fixed depth of 1M reads for all samples (see **Methods**), stratified by SIRV transcript length. R denotes Pearson correlation. **F.** Relative quantification efficiency, computed as number of counts divided by number of raw reads, of four quantification methods for Kinnex lrRNA-seq and Illumina, stratified by the downsampled number of input reads. **G.** Number of significant DTE calls that overlap with at least one other method at the same depth of four quantification methods for Kinnex lrRNA-seq and Illumina, stratified by the downsampled number of input reads. R denotes Pearson correlation.

Next, we proceeded to evaluate transcript quantification performance across three tasks: Differential Transcript Expression (DTE), fold-change estimation, and absolute quantification. First, we asked how well each tool detects Differential Transcript Expression (DTE) between the SIRV E1 and E2 mixes. Isoquant and lr-kallisto performed best, with accuracy comparable to Illumina (Figure 2B). Oarfish and Bambu showed lower DTE accuracy, but this was largely due to an outlier transcript (SIRV303), which was challenging to resolve because of its sequence similarity to other SIRVs (Figure 2C, top). The SIRV303 transcript is fully contained within the SIRV301 transcript. While the expression of SIRV303 stayed constant between the SIRV mixes used for days 0 and 5, the expression of SIRV301 increased 16-fold. Consequently, many of our tested long-read methods attributed most or even all reads of the SIRV303 transcript to the SIRV301 transcript, given the lack of unique alignments to the SIRV303 transcript (Figure 2C, bottom).

Second, we examined the accuracy of the resulting fold-change estimates. Oarfish and Bambu were again affected by the SIRV303 outlier, and Salmon showed consistently poor performance across all metrics (Figure 2D). The results of the remaining methods were concordant with the DTE results.

Third, we compared absolute abundance estimates (log-CPM) to ground truth SIRV concentrations. All tools showed moderate agreement (Pearson r~0.70 overall), while Illumina displayed higher accuracy (r~0.88 overall) (Figure 2E). We also observed a moderate bias across all long-read methods against shorter transcripts (less than 1.25 kb), likely reflecting a limitation in Kinnex library preparation, which we believe to be related to the ratio of SPRI beads used for cleanup during the cDNA synthesis, having picked bead concentrations that would skew towards longer transcripts (Figure 2D, E).

While SIRV spike-ins provide a ground truth, they do not accurately reflect the full complexity of the human transcriptome, so we next assessed tool performance for endogenous transcript quantifications against the full GENCODE annotation. Since a ground truth of endogenous transcripts is not available, as knowledge of the transcriptome is rapidly evolving, we ranked the five tools on two proxy metrics: relative quantification efficiency (fraction of counts over inputted raw reads) and reproducibility of Differential Transcript Expression (DTE) calls across tools (both Kinnex and Illumina Salmon analyses). The relative quantification efficiency measures how efficiently a method turns raw reads into counts. Assuming equally accurate quantification results between methods, this directly correlates with total counts and thus power. The number of significant DTE calls aims to measure power in downstream tasks directly. Since our dataset does not have ground truth for DTE, we aim to remove false positive calls by only counting calls that are shared between two quantification methods (either within Kinnex or between Kinnex and Illumina).

For these metrics, Oarfish outperformed all other tools, including Illumina, achieving strong efficiency and the highest number of reproducible DTE calls (Figure 2F–G). Together with its favorable run-time/memory profile and solid SIRV performance (the only primary deficit being the SIRV303 outlier), we selected Oarfish for all downstream Kinnex analyses.

### Kinnex Outperforms Illumina for DTE by Resolving Transcript-Level Read Ambiguity

Using Oarfish for Kinnex and Salmon for Illumina, we compared gene-level differential expression (DGE) between platforms. Calls were highly concordant between Illumina and Kinnex across matched read depths (Figure 3A), as expected, and previously reported for other long-read technologies [6].

**Figure 3:**
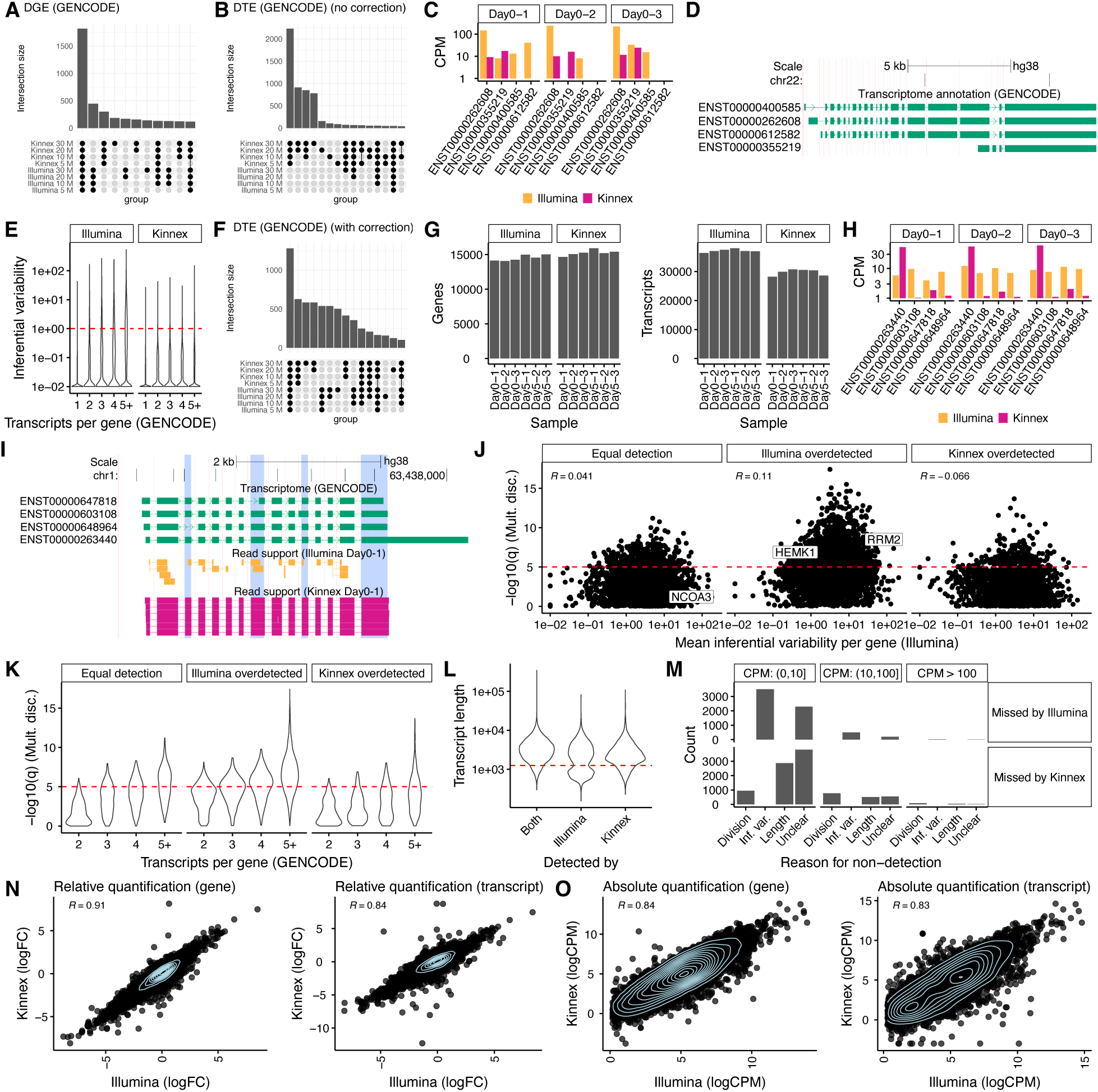
Quantification comparison between Kinnex and Illumina. **A.** Overlap of DGE calls between Day 0 and Day 5 based on Oarfish Kinnex and Salmon Illumina when downsampling GENCODE-aligned data to a fixed read depth for each replicate. **B**. Overlap of DTE calls between Day 0 and Day 5 based on Oarfish Kinnex lrRNA-seq and Salmon Illumina RNA-seq when downsampling GENCODE-aligned data to a fixed read depth for each replicate. **C.** Example of a gene with a ‘flip’, characteristic of transcripts with high inferential variability in Illumina, where Illumina assigns moderate CPM values to a transcript in one replicate, while for another replicate, that transcript has close to zero CPM. **D.** Transcript-model corresponding to the ‘flip’ example (**C**). **E.** Inferential variability (see **Methods**) for GENCODE transcripts according to the number of transcripts per gene, stratified by technology. The y-axis indicates mean inferential relative variance across samples. Per sample, the inferential relative variance measures overdispersion; a value of 100 would indicate that the inferential sample variance is 101 times the inferential sample mean, indicating extreme overdispersion. **F.** Overlap of DTE calls between Day 0 and Day 5 based on Oarfish Kinnex lrRNA-seq and Salmon Illumina RNA-seq when downsampling GENCODE-aligned data to a fixed read depth for each replicate, with Illumina counts corrected for inferential variability (see **Methods**) [11]. **G.** Number of detected transcripts and genes for each replicate of Day 0 and Day 5 when downsampling to 30M reads, stratified by technology. **H.** Example of a ‘division’, where Illumina evenly divides count mass between transcripts with high sequence similarity, whereas Kinnex assigns most count mass to a singular transcript. **I.** Transcript-model corresponding to the ‘division’ example (**H**), with read support for both Illumina and Kinnex, highlighting that both technologies primarily show read support for the ENST00000263440 transcript. **J.** Mean inferential variability per gene in Illumina by the -log10(q) value of testing for multinomial discordance between Illumina and Kinnex, stratified by whether the two platforms detected an equal number of transcripts (equal detection) for the gene in question, or Illumina detected more transcripts (Illumina overdetected) or Kinnex detected more transcripts (Kinnex overdetected). **K.** Number of transcripts per gene by the -log10(q) value of testing for multinomial discordance between Illumina and Kinnex, stratified by whether the two platforms detected an equal number of transcripts (equal detection) for the gene in question, or Illumina detected more transcripts (Illumina overdetected) or Kinnex detected more transcripts (Kinnex overdetected). **L.** Distribution of transcript length for transcripts that are detected by both technologies and transcripts detected only by Kinnex or Illumina, highlighting that Kinnex shows a clear length bias for transcripts shorter than 1,250 nts. **M.** Number of transcripts missed by each technology and possible reasons for their non-detection, stratified by average CPM in the technology in which they were detected. **N.** Relative quantification (log-fold change) concordance between Kinnex and Illumina on the gene and transcript level, for the first replicate, when filtered for transcripts and genes that show low inferential variability in Illumina and are detected by both platforms. **O.** Absolute quantification (log CPM) concordance between Kinnex and Illumina on the gene and transcript level, for the first replicate, when filtered for transcripts and genes that show low inferential variability in Illumina and are detected by both platforms. Unless otherwise noted, Kinnex was downsampled to 30M reads and quantified against GENCODE V45 using Oarfish. Illumina was also downsampled to 30M reads per sample and quantified against the same transcriptome using Salmon (see **Methods**). R denotes Pearson correlation.

Next, we compared DTE calls between Kinnex and Illumina. Kinnex reported many more significant DTE events than Illumina (Figure 3B), consistent with previous work comparing other long-read technologies with Illumina [11]. To explain this discrepancy, we manually inspected several genes in which Kinnex and Illumina quantification differed. In such cases, there was a tendency for the associated genes to express complex transcripts with highly similar sequences. For example, the *CECR2* gene, has four transcripts of highly similar sequence differing only by short exons and alternative acceptor sites (Figure 3C-D). For many such cases, Illumina quantifications exhibited replicate-to-replicate fluctuations in abundances, sometimes dramatically assigning abundances that alternated between near-zero and mid-range values across replicates (”transcript flips”). Kinnex data, on the other hand, did not suffer from this problem, rather assigning expression to one or multiple dominant transcripts in a consistent manner (Figure 3C-D).

To test whether the pattern observed in *CECR2* was an isolated case or a transcriptome-wide phenomenon, we quantified transcript measurement instability across the Kinnex and Illumina results using inferential variability [24]. Consistent with previous findings, we found that transcripts quantified by Illumina short reads exhibited higher inferential variability, as compared to those quantified by long-read technologies such as Kinnex, in our case [10, 11] (Figure 3E). This was particularly true for genes with a higher number of annotated transcripts (Figure 3E).

The consequence of high inferential variability in Illumina data is that it leads to inflated dispersion estimates, ultimately diminishing statistical power to detect true DTE events [11]. This reduced power explains Illumina’s initially small number of DTE calls. Indeed, transcripts with high inferential variability were disproportionately represented among those DTEs missed by Illumina but detected by Kinnex (Figure S3A).

### Correcting for Inferential Variability Improves Illumina Quantification Performance

To focus the benchmarking on transcripts that Illumina can quantify reliably, we first corrected Illumina-associated transcript counts for inferential variability, specifically by down-weighting transcripts with highly uncertain quantification [11]. After correction, the overlap between Illumina and Kinnex DTE calls improved substantially, both in the number of events captured and q-value distributions (Figure 3F, Figure S3A). Accordingly, we employed inferential variability correction on all datasets in this study(including Figure 1 [11]), except where variability itself was the subject of investigation (Figure 3C, H, and Figure S3F-H).

### Illumina’s Apparent Transcript Sensitivity Reflects Ambiguity, Not True Transcript Diversity

Next, we evaluated the sensitivity with which each platform could detect transcripts. For each sample replicate, we determined the number of detected genes and transcripts, using 30M downsampled reads (a transcript or gene was counted as detected if it had one CPM or higher in all samples of the respective day, see **Methods**). Kinnex detected a comparable number of genes on average, compared to Illumina (15,253 vs. 14,506 for Illumina), but notably fewer transcripts (29,721 vs. 37,163; Figure 3G).

Upon closer investigation, we noticed examples in which the additional transcripts detected by Illumina may have been spurious. For the *ALG6* gene, for example, Kinnex identified a single dominant transcript (ENST00000263440), whereas Illumina reported expression that was divided more evenly across all four transcripts (Figure 3H), despite limited unique read support for any of these transcripts (Figure 3I). This suggests that transcripts detected by Illumina could be false positives caused by short-read ambiguity rather than genuine transcript diversity.

In these cases, the overall distribution of transcript proportions of genes reported by Illumina would differ from that reported by Kinnex. To quantify these discrepancies, we employed a multinomial discordance test, which assesses differences in transcript usage between platforms (i.e., DTU, see **Methods**). We found that gene-level discordance (q-value) showed no correlation with the mean inferential variability [24]) (Figure 3J). Similarly to what we previously found with inferential variability, multinomial discordance tended to increase with the number of transcripts per gene, especially for genes for which Illumina detected more transcripts than Kinnex (Figure 3K). Despite these overall trends, we still observed genes with high multinomial discordance but low inferential variability, and vice-versa, implying that transcript divisions and inferential noise are related but distinct sources of error (Figure S3B-D). Finally, consistent with the earlier SIRV findings, Kinnex showed a slight length bias that was independent of the transcript-division issue (Figure 3L). Together, our analyses suggest at least two reasons why quantification differed between platforms. First, Illumina’s short-read limitations led to unreliable quantifications for complex genes, manifested either as transcript flips across replicates or transcript division of expression among multiple similar transcripts. Second, Kinnex exhibited reduced detection efficiency for shorter transcripts, as seen in both spike-in controls and endogenous genes. Despite these biases, transcripts detected by only one platform tended to be lower expressed, especially when no clear technical explanation was evident (Figure 3M).

### Kinnex and Illumina Show Strong Concordance for Reliably Detected Transcripts

Having established the sources of platform-specific transcript detection differences, we then evaluated how well Illumina and Kinnex quantifications agreed for the subset of transcripts reliably detected by both platforms (see **Methods**). This left 8,797 transcripts and 12,197 genes for comparison (note that the lower number of transcripts than genes is due to filtering for inferential variability.). Overall, the two platforms agreed closely. Relative quantification (fold change estimates) was highly correlated with Pearson correlation values exceeding 0.9 at the gene level and approaching 0.9 at the transcript level (Figure 3N). Absolute quantifications (logcounts per million (CPM) values) were somewhat less correlated, but still exceeded values of 0.8 at both the gene and transcript level (Figure 3O). Taken together, within this unbiased subset, Kinnex quantified transcript expression as reliably as Illumina, and did so consistently across the range of transcript lengths (Figure S3E-F).

### Kinnex’s quantification performance remains good at approximately base-normalized depths

Lastly, we investigated whether the good performance of Kinnex data was solely due to the increase in bases relative to Illumina, at equal depth (Figure 1D). For this, we repeated both the transcript discovery experiment on the SIRVs and the quantification comparisons between Illumina and Kinnex. While the median read lengths of Kinnex are ~5.2 times longer than Illumina, we performed the experiments at one tenth of the depth for the SIRV data and one sixth of the depth for Illumina data. This yielded 0.25 M reads for Kinnex and 2.5 M for Illumina for the SIRVs and 5 M reads for Kinnex and 30 M reads for Illumina for GENCODE. Overall, results were concordant with higher read numbers. In particular, Kinnex continued to outperform Illumina in transcript discovery (Figure S3G), and still achieved comparable correlation numbers in both absolute and relative quantification when compared to Illumina at a higher depth (Figure S3H-I). We believe the benefit of Kinnex, and long-read data more broadly, to lie in the lowering of inferential variability, relative to Illumina (Figure 3E). This comes at a cost, given that long-reads are frequently sequenced at a lower depth than Illumina, which may especially affect downstream tasks such as DTE. Despite this, we did not find this to be the case here, given that Kinnex continued to perform well in DTE and DGE, even at a sixth of the depth of Illumina (Figure 3A, F).

## Discussion

Overall, our results provide evidence of strong transcript-level quantification performance for Kinnex. Similarly to other long-read technologies, Kinnex largely seemed to avoid problems of inferential variability and detected more DTE events than Illumina. Even with extensive correction and filtering, Illumina suffered from issues likely caused by short read mapping uncertainties, leading to phenomena such as transcript fluctuations and “flips” as well as artifactual “division” of expression across transcripts, which may erode DTE power. For transcripts reliably quantified across both platforms, Kinnex and Illumina showed high concordance, supporting Kinnex as an effective alternative for transcript-level studies. Nonetheless, Kinnex exhibited some limitations, particularly showing a bias against shorter transcripts and exhibiting a slightly lower performance in absolute compared to relative quantification. In addition, Kinnex and similar lrRNA-seq technologies are typically more expensive than Illumina for the same sequencing depth, though rapid improvements in chemistry and instrumentation are reducing this gap.

Our work extends several recent works focused on evaluating the quantification and transcript discovery capability of lrRNA-seq methods [4–7]. In contrast to previous efforts, we introduce a large-scale Kinnex dataset, sample-matched with Illumina, which newly enables the benchmarking of PacBio data at a similar depth to ONT and Illumina. In addition, we evaluate the quantification performance of Kinnex and highlight its strengths and weaknesses, in particular its good performance at the transcript level through avoiding inferential variability-related pitfalls and its length bias.

These results position Kinnex as a practical, reliable choice for projects that must resolve complex transcript architecture and perform differential expression transcript analyses. Future work should add matched ONT data, deeper coverage datasets across more diverse biological systems, and emerging single-cell long-read methods.

## Methods

### Dataset generation

#### Stem cell culture

Undifferentiated human-induced pluripotent stem cells (hiPSCs, WTC11, NIGMS Repository Number GM25256) were obtained through Coriell via a Materials Transfer Agreement with the University of Virginia. WTC11 cells were thawed from Cryostor (Stem Cell Technologies) and cultured using mTESR Plus Media (Stem Cell Technologies) on Matrigel (Corning) coated plates with media changes following manufacturer recommendations. Cells were cultured to maintain undifferentiated cell populations and passaged when confluent using ReLSR (Stem Cell Technologies) and replated at desired densities per manufacturer guidelines on Matrigel-coated dishes.

#### Stem cell (WTC11) derived primordial endothelial cells

WTC11 cells were seeded with mTESR Plus Media (Stem Cell Technologies) onto 6-well dishes or 10cm Matrigel-coated plates for primordial endothelial cell differentiation as described previously [8]. Twenty-four hours after plating, on Day 0 media was aspirated and replaced with Diff APEL 2 (Stem Cell Technologies) differentiation media containing 5 µm GSK3i (CHIR99021, Reprocell). On Day 1 media was aspirated and replaced with differentiation media containing 50ng/mL essential fibroblast growth factor (bFGF, R and D Systems). On Days 2, 3, and 4 media was aspirated and replaced with differentiation media containing 25ng/mL of bone morphogenetic protein 4 (BMP4, R&D Systems) and 50 ng/mL of vascular endothelial growth factor VEGF (Fisher Scientific). Cells were collected from Days 0-5 of the differentiation protocol using Accutase (ThermoFisher) per manufacturer guidelines and pelleted. Three biological replicates collected on Day 0, with Day 0-1 and Day 0-2 collected from a 10 cm dish and Day 0-3 from a 6-well dish. Three biological replicates were collected as described above for Day 5, with Day 5-1 and Day 5-2 derived from a 10 cm dish and Day 5-3 from a 6-well plate. Two biological replicates were obtained from Day 3 from 10 cm dishes. Days 1, 2, and 4 were all collected from 10 cm dishes. For each sample, RNA was extracted at approximately 1 million cells using an RNeasy (Qiagen) kit, with quality assessment via Bioanalyzer. Spike-In RNA Variants (SIRVs, Lexogen) were reconstituted per manufacture guidelines and added to samples with Day 0 biological replicates receiving E1 SIRVs, Day 5 E2, and all other Days receiving S4. Master mixes of RNA and SIRVs were split into 4 tubes for parallel long-read Pacific Biosciences (PB) Revio sequencing and short-read sequencing on the NovaSeq at 150bp.

#### qPCR gene marker analysis

Transcriptomes were collected from cultured WTC11 cells at every day of the differentiation protocol using the RNeasyPlus Micro Kit (QIAGEN). Resulting cDNA libraries were assessed via qPCR using published PCR primers (Integrated DNA Technologies, based on [8, 25]) and miScript SYBR Green PCR Kit (QIAGEN) to validate gene expression patterns associated with the intermediate states.

#### Short-read RNA-sequencing

Aliquots of total RNA samples created from the master mix preparation as described above, were used as input for sequencing to Kapa RNA HyperProKits for short-read RNA sequencing library preparation. The resulting libraries were sequenced on the NovaSeq at 150bp.

#### Kinnex long-read RNA sequencing library preparation and long-read RNA sequencing run

Aliquots of total RNA samples created from the master mix preparation as described above were used as input for PB Kinnex. From the RNA, cDNA was synthesized using the Iso-Seq Express Kit (PB). Approximately 300ng of cDNA from each sample was barcoded and equally pooled. All replicates in one WTC11 sample were made into one Kinnex full-length library and sequenced on one SMRTCell (i.e., WTC11 Day 0 and WTC11 Day 5 replicates were sequenced in one run).

### Data Analysis

#### Preprocessing

To prepare raw sequencing files for alignment, we performed different preprocessing steps per technology.

##### Illumina

We ran Trim Galore 0.6.2 in paired-end mode with a minimum length of 20 and a minimum quality of 20 using the `--phred33` option.

##### Kinnex

Kinnex data were preprocessed on-instrument. Basecalling was performed using basecaller version 5.0 and otherwise standard parameters. CCS reads were generated using ccs 7.0.0, followed by skera demultiplexing of Kinnex S-reads using skera 1.0.99, lima 2.8.99, and Isoseq refine 3.99.99, at which point reads were ready for alignment.

#### Separating FASTQ into spike-in and non-spike-in data

As an initial step, both for QC and since spike-ins may require different alignment options, we separated all FASTQ files into reads aligning to either one of the artificial spike-in genomes or to the human genome.

##### Alignment for Illumina-generated RNA-sequencing files

Data was aligned to a concatenation of the human genome, the full SIRV genome, and the ERCC genome using STAR 2.7.11b [26]. Afterward, BAM files were separated by whether they aligned to the human or one of the artificial genomes and converted to FASTQ using samtools 2.19.2 [27].

##### Alignment for Kinnex-generated RNA-sequencing files

Data was aligned to a concatenation of the human genome, the full SIRV genome, and the ERCC genome using minimap2 2.28-0 with the `splice:hq -uf` options [28]. Afterward, BAM files were separated by whether they aligned to the human or one of the artificial genomes and converted to FASTQ using samtools 2.19.2.

#### Genome alignment (Kinnex)

Genome alignment was performed using minimap2 2.28-0 with the `splice:hq -uf` options and provided annotated splice sites using `--junc-bed`. For the alignment of SIRV reads, we also applied the `--splice-flank` option.

#### Transcriptome alignment (Kinnex)

Transcriptome alignment was performed using minimap2 2.28-0 with the `map-hifi --eqx` options, allowing for a maximum of 100 secondary alignments by using the `-N 100` option.

#### Genome alignment (Illumina)

Genome alignment was performed using STAR 2.7.11b using the `--outSAMstrandField intronMotif` argument and otherwise default parameters. Indices for STAR were generated using the `--sjdbGTFfile` option to which appropriate transcriptome GTF files were passed. The `--sjdbOverhang` option was set to 150.

#### Transcriptome alignment (Illumina)

Transcriptome alignment was performed using STAR 2.7.11b using default parameters. Indices for STAR were generated using default parameters.

#### References

We used the GRCh38 genome (`GCA 000001405.15 GRCh38 no alt analysis set.fna.gz`) as the human reference genome and the primary assembly of GENCODE V45 as the human reference transcriptome [29]. For the spike-ins, we used the genome and transcriptome corresponding to SIRV set four, including short SIRVs, long SIRVs, and ERCCs.

#### Quantification (Kinnex)

##### Isoquant

Isoquant 3.6.1 was used to quantify genome alignments with precomputed DB files, never requiring a polyA tail, data type `pacbio_ccs`, and otherwise default parameters [21].

##### lr-kallisto

kallisto 0.51.0 was used in conjunction with bustools 0.44.1 to quantify FASTQ files using an index k-mer setting of 63, a bus threshold of 0.8, and PacBio as the platform (`-P PacBio`) [22].

##### Oarfish

Oarfish 0.6.2 was run to quantify transcriptome alignments with model coverage applied, no filters, and otherwise default parameters [23].

##### Bambu

Bambu 3.8.0 was run to quantify genome alignments with default parameters [17].

##### Salmon

Salmon 1.10.3 was run to quantify transcriptome alignments with the `--ont` flag and otherwise default parameters.

#### Quantification (Illumina)

For Illumina data, we used Salmon 1.10.3 to quantify FASTQ files using an index k-mer setting of 31, a fragment length prior mean of 250, a fragment length standard deviation of 25, the `--validateMappings --gcBias --seqBias` flags, and otherwise default parameters [20].

#### Inferential variability correction

For Illumina, we corrected for inferential variability using the `catchSalmon` function in edgeR 4.4.0. Kinnex quantifications were not corrected for inferential variability [11, 30].

#### Gene-level quantifications

Gene-level quantifications were generated by summing all transcript-level quantifications for all transcripts belonging to a gene, according to a particular annotation.

#### Quality control

##### Read count numbers

Read count numbers were calculated using samtools 2.19.2. For aligned data, we counted primary alignments after separation into SIRV and GENCODE data.

##### Read length and number of bases

Read length was calculated using bioawk 1.0 as the number of nucleotides in each read for Kinnex and the sum of nucleotides in each paired-end read for Illumina. Number of bases was calculated as the sum of read lengths for each technology.

##### Number of covered junctions

Covered junctions were calculated as the number of introns of length 20 or longer in the cigar string of the genome alignments of each technology using pysam 0.23.0.

##### Read quality and relative edit distance

Read quality was calculated as the average phred value of all bases of that read for Kinnex or the average phred of all bases in each paired-end read for Illumina using bioawk 1.0.

Relative edit distance was heuristically calculated based on the BAM file as the edit distance from the nM tag divided by the total number of aligned bases for Kinnex and the sum of edit distances divided by the sum of the total number of aligned bases in each paired-end read for Illumina using pysam 0.23.0.

##### Gene body coverage

Gene body coverage was calculated on genome-aligned BAM files using randomly subsampled GENCODE transcripts using RSeQC 5.0.4 [31]. Gene body coverage was calculated per transcript and within size bins. For each size bin, we randomly sampled 2,500 GENCODE transcripts.

##### Fragment size

For Illumina data, fragment size was calculated on transcriptome-aligned BAM files by using the TLEN attribute of the BAM file using pysam 0.23.0.

#### Transcript and gene filtering

Depending on the analysis, we performed different types of transcript or gene filtering. SIRV transcripts were never filtered.

##### Replicability (Figure 1 J-K)

Only transcripts or genes having at least one CPM in at least three samples in both platforms were kept.

##### QC detection (Figure 1P and Figure 2G)

A transcript or gene was considered as detected if it had at least one CPM in the sample in question.

##### Differential analyses

For all differential analyses, we used the filterByExpr function of edgeR 4.4.0 with default parameters.

##### Technology detection (Figure 2J-M)

Transcripts were considered as detected if they had at least one CPM in all three replicates of day zero.

##### Relative quantification concordance (Figure 2N)

Transcripts were kept if they had inferential variability less than five for the Illumina data, and they had at least one CPM in all three replicates of day zero in both technologies. Requirements for genes were identical except that no inferential variability filter was applied.

##### Absolute quantification concordance (Figure 2O)

Transcripts were kept if they had inferential variability less than one for the Illumina data, and they had at least one CPM in all three replicates of day zero in both technologies. Requirements for genes were identical except that no inferential variability filter was applied.

#### Transcript discovery (GENCODE)

Bambu 3.8.0 was run to perform transcript discovery using Kinnex genome alignments, selecting the Bambu-suggested NDR and keeping ISM novel transcripts (`remove.subsetTx = FALSE`).

#### Transcript discovery (SIRVs)

Transcript discovery on the SIRVs was ran on SIRV genome alignments, downsampled to 2.5 M reads for both platforms. Genome alignments were generated without providing the location of existing SIRV splice junctions to the aligner.

Discovery for Illumina was performed using stringtie 2.2.3 with default parameters and without a reference transcriptome. Discovery for Kinnex was performed using Bambu 3.8.0 in de-novo mode, with an NDR of 1.0 and keeping ISM novel transcripts (`remove.subsetTx = FALSE`) [32].

Performance was calculated using gffcompare 0.12.6 by comparing the predicted transcriptomes of both technologies to the ground truth transcriptome of the short SIRVs [33]. Performance metrics were reported at the intron chain level.

#### Annotation of novel transcripts (GENCODE)

Novel transcripts were annotated using SQANTI3 5.1.2. For determining whether a particular novel transcript was coding or non-coding, we ran ORFanage 1.1.0 in `BEST` mode, using GENCODE V45 CDS as a reference, and determined a novel transcript as coding if it was assigned a CDS by ORFanage [2, 18].

#### Inferential variability

We used `fishpond` 2.12.0 for all analyses regarding inferential variability. In particular, we used `computeInfRV` to calculate inferential variability as the mean inferential relative variance of each transcript across samples [24].

#### Quantification of transcript mass division between Illumina and Kinnex

To quantify differences in the division of transcript mass for a particular gene between Illumina and Kinnex, we performed a test for multinomial discordance between Illumina and Kinnex. In particular,DTU was performed with edgeR’s `glmQLFit` and `diffSpliceDGE` across all replicates of day zero of both Illumina and Kinnex, accounting for both technology and replicate. We tested significance on the technology coefficient. Transcript-level q-values were either used directly or aggregated to the gene level using Simes’ adjustment.

#### Definition of categories leading to lack of detection (Figure 2M)

##### Transcripts missed by Kinnex

1. **Length**: Transcripts that were shorter than 1,250 nucleotides
2. **Division**: Transcripts longer than 1,250 nucleotides but having a -log10(q) of multinomial discordance between Illumina and Kinnex greater than five
3. **Unclear**: All other transcripts not detected by Kinnex but detected by Illumina

##### Transcripts missed by Illumina

1. **Inf. var.**: Transcripts with a mean inferential variability value greater than five
2. **Unclear**: All other transcripts not detected by Illumina but detected by Kinnex

#### Downsampling

For optimal comparability in some analyses, we downsampled read FASTQ or alignment BAM files without replacement.

SIRV files were downsampled to 0.25 million, 0.5 million, 1 million, and 2.5 million, while GENCODE files were downsampled to 5, 10, 20, and 30 million, respectively. For the SIRV downsamplings, we performed five downsamplings per target depth with different random seeds, while for GENCODE only one downsampling was performed.

BAM files for genome and transcriptome alignments were downsampled using a script leveraging Rsamtools 2.18.0 that kept all alignments corresponding to a source read. FASTQ files were downsampled using seqtk 1.4, ensuring contiguity for paired-end Illumina reads.

#### Differential analyses (GENCODE)

Differential Gene Expression (DGE) was performed between all replicates of Day 0 and Day 5, using edgeR’s `glmQLFit`, accounting for both day and replicate. We tested significance on the day coefficient and called genes as significant that had a q-value less than 0.01.

DTE was performed between all replicates of Day 0 and Day 5, using edgeR’s `glmQLFit`, accounting for both day and replicate. We tested significance on the day coefficient and called transcripts as significant that had a q-value less than 0.01.

DTU was performed across all replicates across the timecourse (excepting Day2-1) using a cubic spline with edgeR’s `glmQLFit` and `diffSpliceDGE`, accounting for both day and replicate. We tested significance on all spline coefficients jointly and called genes as significant that had a q-value less than 0.01, based on Simes adjustment.

#### Differential analyses (SIRVs)

We performed DTE on the SIRV datasets between all replicates of Day 0 and Day 5, corresponding to SIRV mixes E1 and E2, respectively. In particular, we used the true log-fold changes between SIRV mixes E1 and E2 as the ground-truth, yielding either 0 (no change between mixes), −1 (decrease from E1 to E2), or 1 (increase from E1 to E2).

We then performed DTE using edgeR’s `glmQLFit`, with a design matrix accounting for both the mix and the replicate, testing the significance of the mix coefficient. Predictions were extracted using limma’s `decideTests`, using an FDR cut-off of 1% and no log-fold change cutoff [34]. The FDR was calculated as the percentage of SIRVs found as −1 or 1 that were either 0 or the opposing sign in the ground-truth (i.e., false discoveries) over all discoveries (i.e., all SIRVs that had a non-zero value assigned). The TPR was calculated as the percentage of SIRVs that had a ground-truth non-zero value, were deemed significant, and had the same sign as the ground-truth over all ground-truth non-zero values.

All quantifications were downsampled to a depth of one million reads for fairness. FDR and TPR values correspond to mean values calculated across all five downsamplings.

#### Relative quantification efficiency

Relative quantification efficiency was assessed for each quantification method by calculating the average proportion of quantified counts to raw input reads at several downsampled depths for the GENCODE data (5M, 10M, 20M, 30M).

#### Definition of significant DTE calls (Figure S2G)

We considered any call that had an FDR less than 0.01 and was called by at least one other quantification method at the same depth as significant.

#### Computational requirements

Computational requirements were measured using the Snakemake `benchmark` functionality and are reported over each of the three replicates for days zero and five. Since computational requirement variability was minimal for reruns, we do not report variability within replicate and day. All methods were run with access to 12 cores.

#### Reproducibility

All bioinformatics experiments were performed on an Ubuntu 22.04.5 LTS server with AMD EPYC 7742 CPUs. Experiments were run with Snakemake 8.2.3 within rule-appropriate mamba environments within Singularity containers [35]. All experiments are reproducible up to numerical stability via standard Snakemake commands documented on our Github (see **Data Availability**).

All analysis was either performed in respective command line tools or in R 4.4.3 [36]. Figures were generated using ggplot2 3.5.2 and ggpubfigs 1.1.0 [37, 38].

## Data Availability

For access to the raw sequencing files for both Illumina and Kinnex, see [University of Virginia (UVA)](https://portal.ithriv.org/%23/public_commons/project/25d25b41-87ca-4e41-9573-1ee9375daf1a/dataset/5bcbeb1e-a0e6-4dd3-91ed-107eba1d3061), [Integrated Translational Health Research Institute of Virginia (iTHRIV)](https://portal.ithriv.org/%23/public_commons/project/25d25b41-87ca-4e41-9573-1ee9375daf1a/dataset/5bcbeb1e-a0e6-4dd3-91ed-107eba1d3061). Researchers who wish to gain access to the source data can contact the authors to initiate a data request. Before sharing, both groups will seek approval from the NIGMS repository by completing the Statement of Research Intent forms. Once approved, a contract can be initiated between institutions for the transfer and reuse of the source data in this study.

Count tables, separated into spike-in and non-spike-in data for the GENCODE reference transcriptome, and the GENCODE reference transcriptome augmented with Kinnex lrRNA-seq-derived novel transcripts for both Illumina and four quantification methods for PB Kinnex (**Methods**), along with related paper outputs have been deposited to [Zenodo](https://doi.org/10.5281/zenodo.15502054). Other results that either required controlled access deposition or were too large for deposition via Zenodo are available upon demand.

All code to reproduce experiments, figures, and tables is available on [Github](https://github.com/dnwissel/kinnex-data).

## Acknowledgements

We would like to acknowledge Pacific Biosciences of California, Inc., for providing sequencing of PB Kinnex lrRNA-seq on the Revio free of charge. Early results based on the dataset introduced here were presented in a webinar organised by Pacific Biosciences of California, Inc. We are grateful to webinar attendees for their helpful feedback and discussions. Figure 1A was created with Biorender.com (publication license available on demand). This work was supported by a Swiss National Science Foundation (SNSF) project grant 204869 to M.D.R., the Robert M. Berne Cardiovascular Research Center Training Program (T32HL007284), the Wagner Fellowship (UVA) to M.M.M, and a National Institute of General Medical Sciences (NIGMS), National Institutes of Health (NIH) grant R35GM142647 to G.M.S.

## Author Information

These authors contributed equally: David Wissel, Madison M. Mehlferber.

## Contributions

G.M.S. and M.D.R. conceived of the study design, and M.M. implemented the cell-based experiments. M.M. and V.P. performed experimental work and cell collection. M.M. and G.M.S. coordinated sequencing of the Illumina RNA-seq samples. E.T. coordinated the sequencing of PB Kinnex lrRNA-seq samples. D.W., M.M., and K.N. wrote the analysis code. D.W. wrote the Snakemake workflow. D.W. and M.M. analyzed the results. M.M. and D.W. drafted the original manuscript. E.T., M.D.R., and G.M.S. co-supervised the work. All authors reviewed, edited, and approved the final manuscript.

## Declarations

E.T. is employed by and holds stock in Pacific Biosciences of California, Inc. G.M.S. serves on the scientific advisory board of Quantum-Si Incorporated and holds stock in Quantum-Si Incorporated. Pacific Biosciences of California, Inc. ran sequencing of PB Kinnex lrRNA-seq on the Revio free of charge.

All remaining authors declare no competing interests.

---

**Figure S1:**
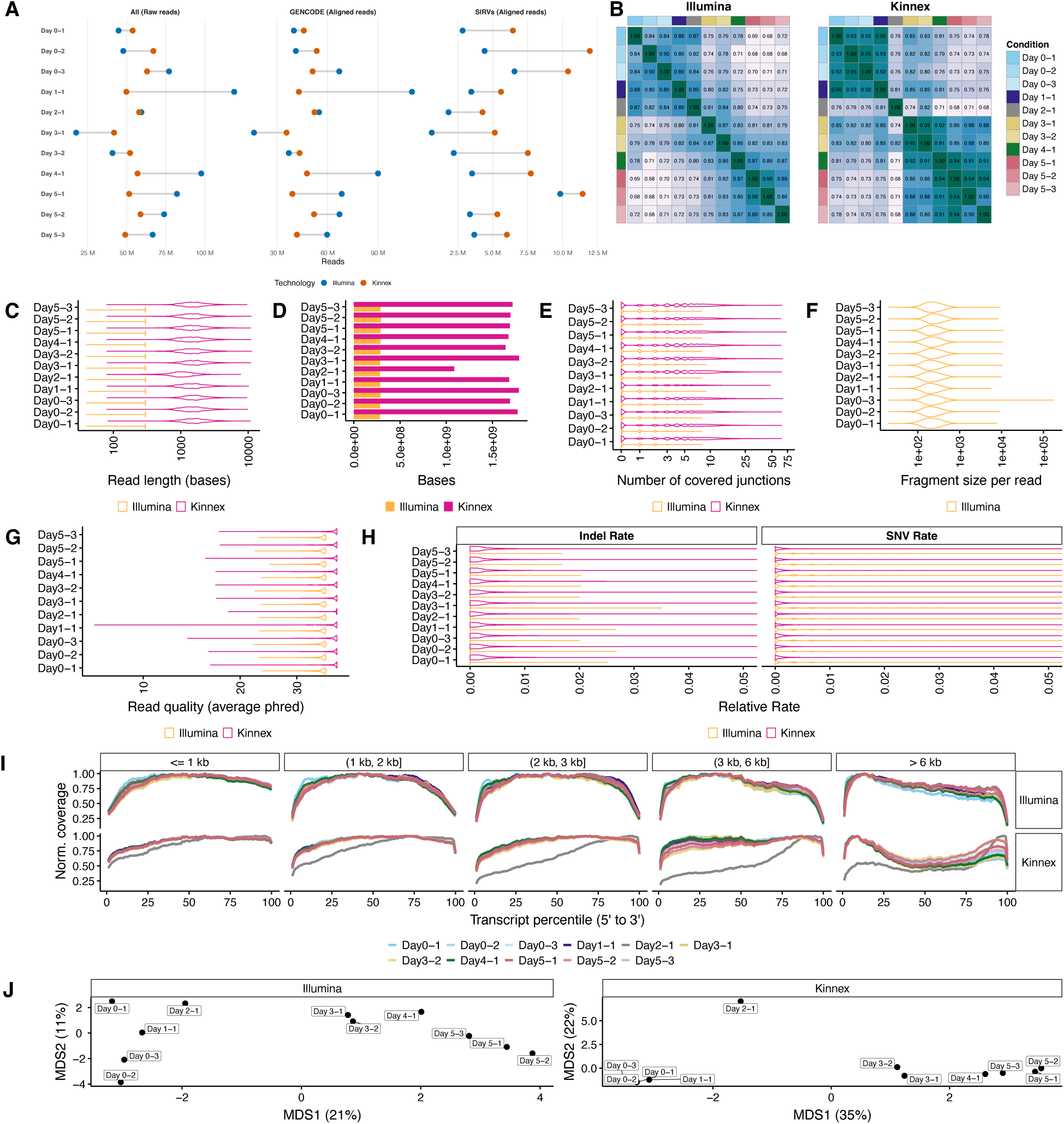
Quality control of all samples in our dataset highlights quality issues in Day2-1 for Kinnex. **A.** Number of raw and aligned reads per technology, stratified by their source across all samples. **B.** Spearman correlation heatmap of transcript-level quantification across all samples, highlighting sample-similarities across the full differentiation for each technology and quality issues with Day 2-1 for Kinnex. **C.** Read lengths of 1M randomly sampled reads aligning to the human genome, stratified by technology for all samples, highlighting quality issues with Day2-1 for Kinnex. For Illumina, the length was calculated across both ends. **D.** Number of base pairs sequenced from 1M randomly sampled reads collected for each technology (same reads as **C**). **E.** Number of covered junctions per read from 1M randomly sampled reads collected for each technology (same reads as **C**). **F.** Fragment size distribution of 1M randomly sampled reads collected for each technology (same reads as **C**). **G.** Average base quality of 1M randomly sampled reads aligning to the human genome, stratified by technology for all days (same reads as **C**). Average base quality for Illumina was determined from both ends. **H.** Average edit distance per read, stratified by indel vs SNV errors, of 1M randomly sampled reads collected for each technology (same reads as **C**). **I.** Normalized coverage of reads aligning to 2,500 randomly sampled GENCODE transcripts each across gene body percentiles, stratified by technology and transcript length, across all samples, highlighting quality issues with Kinnex Day2-1. **J.** Transcript-level quantification-based MDS plots highlighting quality issues for Day2-1 for Kinnex, stratified by technology.

**Figure S2:**
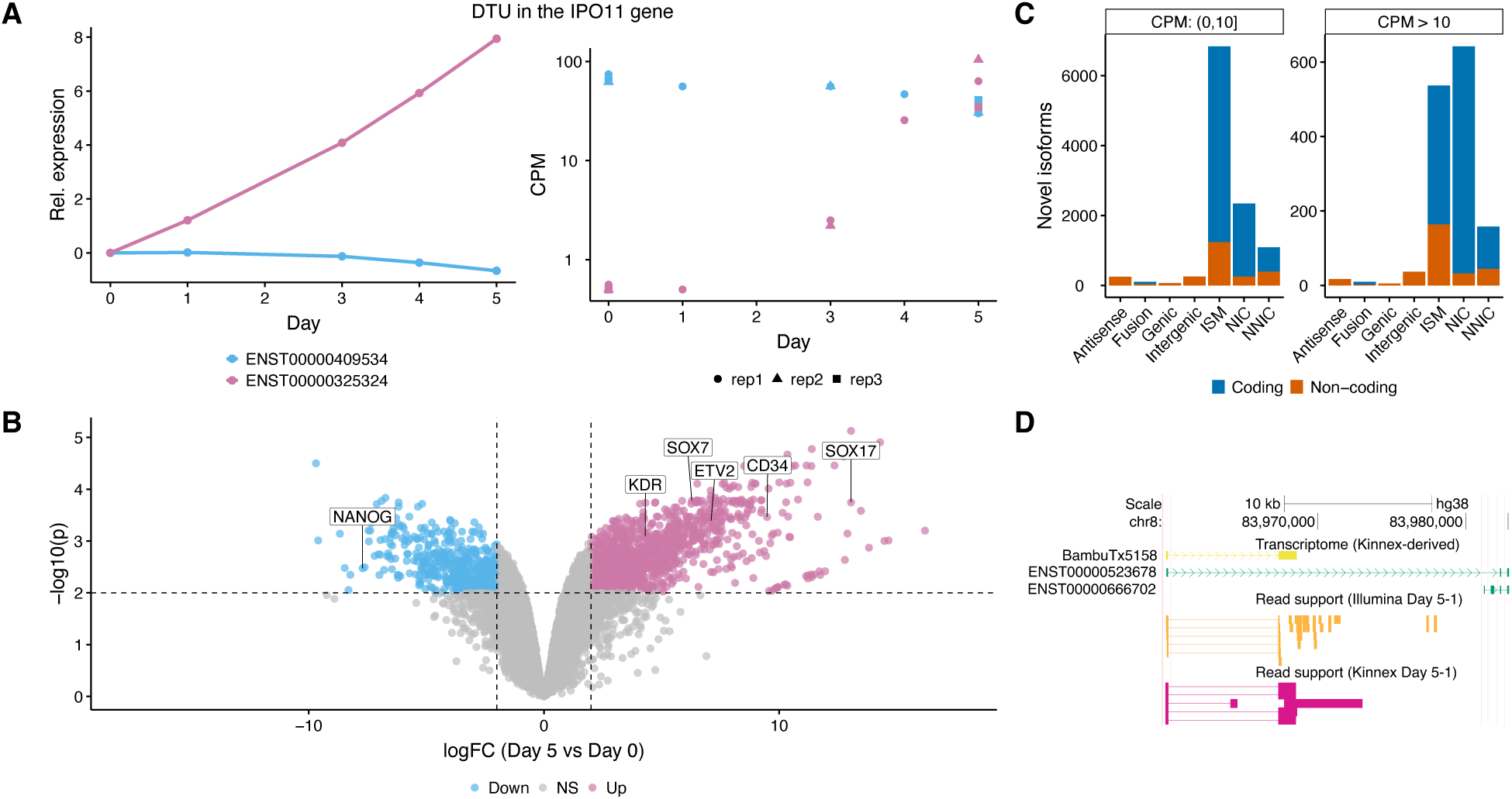
Biological exploration on the iPSC-EC differentiation dataset reveals differential splicing, differential expression and a moderate amount of novel isoforms. **A.** Estimated relative (left) and absolute observed (right) expression of two transcripts belonging to the *IPO11* gene across the differentiation. Relative expression estimated using a cubic spline with two degrees of freedom (see **Methods**). **B.** Volcano plot of Differential Gene Expression between Day 0 and Day 5, highlighting genes specific to pluripotency (light blue) and primordial endothelial cells (pink). **C.** Number of novel transcripts in each category as annotated by SQANTI3, stratified by mean CPM in the two samples with the highest CPM for a particular transcript, colored by coding status as predicted by ORFanage. **D.** Browser track highlighting an example of a relatively confidently identified novel transcript that has read support in both Kinnex and Illumina data.

**Figure S3:**
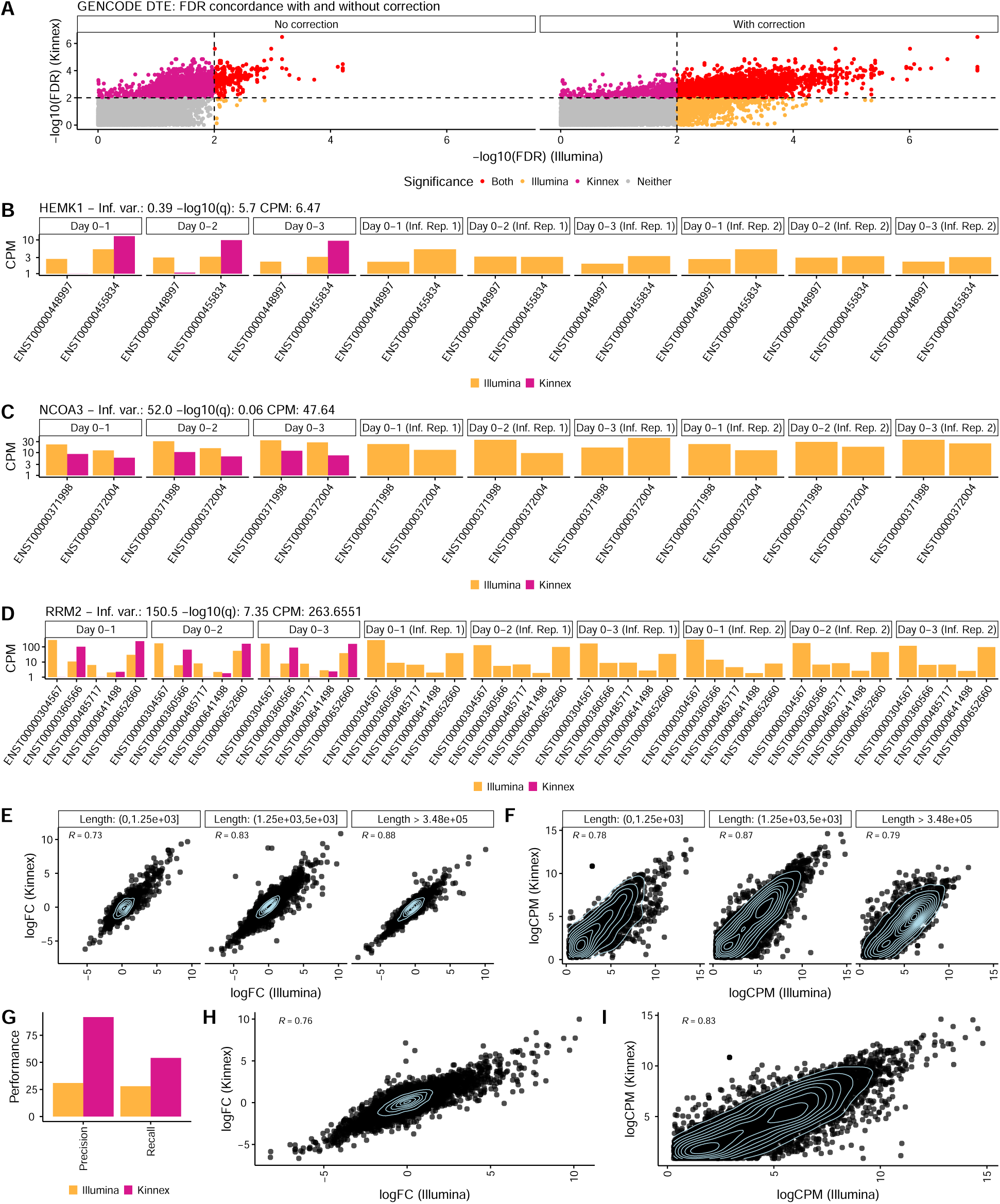
Performance of Kinnex relative to Illumina on DTE, and relative and absolute quantification, including when downsampled to approximately base-normalized depth. **A.** Concordance in per-transcript DTE q-values between Illumina and Kinnex with and without applying an inferential variability correction for Illumina [11] (see **Methods**). Only transcripts that were not filtered for all technologies (Kinnex, Illumina without correction, Illumina with correction) are shown. **B.** Exemplary quantification of the three Day 0 replicates for Illumina and Kinnex and two inferential Day 0 replicates for Illumina on the *HEMK1* gene. **C.** Exemplary quantification of the three Day 0 replicates for Illumina and Kinnex and two inferential Day 0 replicates for Illumina on the *NCOA3* gene. **D.** Exemplary quantification of the three Day 0 replicates for Illumina and Kinnex and two inferential Day 0 replicates for Illumina on the *RRM2* gene. R denotes Pearson correlation. **E.** Concordance of the first replicate per-transcript log-fold changes between Kinnex and Illumina stratified by length in nts. **F.** Concordance of the first replicate of Day 0 per-transcript log CPM values between Kinnex and Illumina, stratified by length in nts. For fairness, in panels E-F, both technologies were downsampled to 30M reads. **G.** Precision and recall of de-novo transcript discovery on three Day 0 (E1 mix) SIRV replicates by technology, when run on approximately base-normalized data. Illumina was run on a downsampled read set consisting of 2.5 M reads while Kinnnex was run on a downsampled read set of 0.25 M reads. **H.** Concordance of the first replicate per-transcript log-fold changes between Kinnex and Illumina when run on approximately base-normalized data. Illumina was run on 30 M downsampled reads, while Kinnex was run on 5 M. **I.** Concordance of the first replicate per-transcript log CPM values between Kinnex and Illumina. Illumina was run on 30 M downsampled reads, while Kinnex was run on 5 M.

## References

[1] Monzó, C., Liu, T., Conesa, A.: Transcriptomics in the era of long-read sequencing. Nature Reviews Genetics, 1–21 (2025)

[2] Pardo-Palacios, F.J., Arzalluz-Luque, A., Kondratova, L., Salguero, P., Mestre-Tomás, J., Amorín, R., Estevan-Morió, E., Liu, T., Nanni, A., McIntyre, L., et al.: Sqanti3: curation of long-read transcriptomes for accurate identification of known and novel isoforms. Nature methods 21(5), 793–797 (2024)

[3] Al’Khafaji, A.M., Smith, J.T., Garimella, K.V., Babadi, M., Popic, V., Sade-Feldman, M., Gatzen, M., Sarkizova, S., Schwartz, M.A., Blaum, E.M., et al.: High-throughput rna isoform sequencing using programmed cdna concatenation. Nature biotechnology 42(4), 582–586 (2024)

[4] Pardo-Palacios, F.J., Wang, D., Reese, F., Diekhans, M., Carbonell-Sala, S., Williams, B., Loveland, J.E., De María, M., Adams, M.S., Balderrama-Gutierrez, G., et al.: Systematic assessment of long-read rna-seq methods for transcript identification and quantification. Nature methods 21(7), 1349–1363 (2024)

[5] Chen, Y., Davidson, N.M., Wan, Y.K., Yao, F., Su, Y., Gamaarachchi, H., Sim, A., Patel, H., Low, H.M., Hendra, C., et al.: A systematic benchmark of nanopore long-read rna sequencing for transcript-level analysis in human cell lines. Nature methods, 1–12 (2025)

[6] Dong, X., Du, M.R., Gouil, Q., Tian, L., Jabbari, J.S., Bowden, R., Baldoni, P.L., Chen, Y., Smyth, G.K., Amarasinghe, S.L., et al.: Benchmarking long-read rna-sequencing analysis tools using in silico mixtures. Nature Methods 20(11), 1810–1821 (2023)

[7] Su, Y., Yu, Z., Jin, S., Ai, Z., Yuan, R., Chen, X., Xue, Z., Guo, Y., Chen, D., Liang, H., et al.: Comprehensive assessment of mrna isoform detection methods for long-read sequencing data. Nature Communications 15(1), 3972 (2024)

[8] Nelson, E.A., Qiu, J., Chavkin, N.W., Hirschi, K.K.: Directed differentiation of hemogenic endothelial cells from human pluripotent stem cells. J. Vis. Exp. (169) (2021)

[9] Soneson, C., Yao, Y., Bratus-Neuenschwander, A., Patrignani, A., Robinson, M.D., Hussain, S.: A comprehensive examination of nanopore native rna sequencing for characterization of complex transcriptomes. Nature communications 10(1), 3359 (2019)

[10] Dong, X., Du, M.R.M., Gouil, Q., Tian, L., Jabbari, J.S., Bowden, R., Baldoni, P.L., Chen, Y., Smyth, G.K., Amarasinghe, S.L., Law, C.W., Ritchie, M.E.: Benchmarking long-read RNA-sequencing analysis tools using in silico mixtures. Nat. Methods 20(11), 1810–1821 (2023)

[11] Baldoni, P.L., Chen, Y., Hediyeh-Zadeh, S., Liao, Y., Dong, X., Ritchie, M.E., Shi, W., Smyth, G.K.: Dividing out quantification uncertainty allows efficient assessment of differential transcript expression with edger. Nucleic Acids Research 52(3), 13–13 (2024)

[12] Zhang, C., Li, H., Wang, S.: Common gene signatures and molecular mechanisms of diabetic nephropathy and metabolic syndrome. Front. Public Health 11, 1150122 (2023)

[13] Romero, M.F., Chen, A.-P., Parker, M.D., Boron, W.F.: The slc4 family of bicarbonate hco transporters. Mol. Aspects Med. 34(2-3), 159–182 (2013)

[14] Liao, Z., Cantor, J.M.: Endothelial cells require CD98 for efficient angiogenesis-brief report. Arterioscler. Thromb. Vasc. Biol. 36(11), 2163–2166 (2016)

[15] Nichol, D., Stuhlmann, H.: EGFL7: a unique angiogenic signaling factor in vascular development and disease. Blood 119(6), 1345–1352 (2012)

[16] Yeh, Y.-T., Hur, S.S., Chang, J., Wang, K.-C., Chiu, J.-J., Li, Y.-S., Chien, S.: Matrix stiffness regulates endothelial cell proliferation through septin 9. PLoS One 7(10), 46889 (2012)

[17] Chen, Y., Sim, A., Wan, Y.K., Yeo, K., Lee, J.J.X., Ling, M.H., Love, M.I., Göke, J.: Context-aware transcript quantification from long-read rna-seq data with bambu. Nature methods 20(8), 1187–1195 (2023)

[18] Varabyou, A., Erdogdu, B., Salzberg, S.L., Pertea, M.: Investigating open reading frames in known and novel transcripts using orfanage. Nature computational science 3(8), 700–708 (2023)

[19] Amarasinghe, S.L., Ritchie, M.E., Gouil, Q.: long-read-tools. org: an interactive catalogue of analysis methods for long-read sequencing data. GigaScience 10(2), 003 (2021)

[20] Patro, R., Duggal, G., Love, M.I., Irizarry, R.A., Kingsford, C.: Salmon provides fast and bias-aware quantification of transcript expression. Nature methods 14(4), 417–419 (2017)

[21] Prjibelski, A.D., Mikheenko, A., Joglekar, A., Smetanin, A., Jarroux, J., Lapidus, A.L., Tilgner, H.U.: Accurate isoform discovery with isoquant using long reads. Nature Biotechnology 41(7), 915–918 (2023)

[22] Loving, R.K., Sullivan, D.K., Booeshagi, A.S., Reese, F., Rebboah, E., Sakr, J., Rezaie, N., Liang, H.Y., Filimban, G., Kawauchi, S., et al.: Long-read sequencing transcriptome quantification with lr-kallisto. bioRxiv, 2024–07 (2025)

[23] Jousheghani, Z.Z., Patro, R.: Oarfish: Enhanced probabilistic modeling leads to improved accuracy in long read transcriptome quantification. bioRxiv (2024)

[24] Zhu, A., Srivastava, A., Ibrahim, J.G., Patro, R., Love, M.I.: Nonparametric expression analysis using inferential replicate counts. Nucleic Acids Research 47(18), 105–105 (2019)

[25] Qiu, J., Nordling, S., Vasavada, H.H., Butcher, E.C., Hirschi, K.K.: Retinoic acid promotes endothelial cell cycle early G1 state to enable human hemogenic endothelial cell specification. Cell Rep. 33(9), 108465 (2020)

[26] Dobin, A., Davis, C.A., Schlesinger, F., Drenkow, J., Zaleski, C., Jha, S., Batut, P., Chaisson, M., Gingeras, T.R.: Star: ultrafast universal rna-seq aligner. Bioinformatics 29(1), 15–21 (2013)

[27] Li, H., Handsaker, B., Wysoker, A., Fennell, T., Ruan, J., Homer, N., Marth, G., Abecasis, G., Durbin, R., Subgroup, .G.P.D.P.: The sequence alignment/map format and samtools. bioinformatics 25(16), 2078–2079 (2009)

[28] Li, H.: Minimap2: pairwise alignment for nucleotide sequences. Bioinformatics 34(18), 3094–3100 (2018)

[29] Frankish, A., Carbonell-Sala, S., Diekhans, M., Jungreis, I., Loveland, J.E., Mudge, J.M., Sisu, C., Wright, J.C., Arnan, C., Barnes, I., et al.: Gencode: reference annotation for the human and mouse genomes in 2023. Nucleic acids research 51(D1), 942–949 (2023)

[30] Robinson, M.D., McCarthy, D.J., Smyth, G.K.: edger: a bioconductor package for differential expression analysis of digital gene expression data. bioinformatics 26(1), 139–140 (2010)

[31] Wang, L., Wang, S., Li, W.: Rseqc: quality control of rna-seq experiments. Bioinformatics 28(16), 2184–2185 (2012)

[32] Pertea, M., Pertea, G.M., Antonescu, C.M., Chang, T.-C., Mendell, J.T., Salzberg, S.L.: Stringtie enables improved reconstruction of a transcriptome from rna-seq reads. Nature biotechnology 33(3), 290–295 (2015)

[33] Pertea, G., Pertea, M.: Gff utilities: Gffread and gffcompare. F1000Research 9, (2020)

[34] Ritchie, M.E., Phipson, B., Wu, D., Hu, Y., Law, C.W., Shi, W., Smyth, G.K.: limma powers differential expression analyses for rna-sequencing and microarray studies. Nucleic acids research 43(7), 47–47 (2015)

[35] Köster, J., Rahmann, S.: Snakemake—a scalable bioinformatics workflow engine. Bioinformatics 28(19), 2520–2522 (2012)

[36] R Core Team: R: A Language and Environment for Statistical Computing. R Foundation for Statistical Computing, Vienna, Austria (2025). R Foundation for Statistical Computing. https://www.R-project.org/

[37] Wickham, H.: Ggplot2: Elegant Graphics for Data Analysis. Springer,(2016). https://ggplot2.tidyverse.org

[38] Steenwyk, J.L., Rokas, A.: ggpubfigs: colorblind-friendly color palettes and ggplot2 graphic system extensions for publication-quality scientific figures. Microbiology Resource Announcements 10(44), 10–1128 (2021)

